# PHRED-1 is a divergent neurexin-1 homolog that organizes muscle fibers and patterns organs during regeneration

**DOI:** 10.1101/131789

**Authors:** Carolyn E. Adler, Alejandro Sánchez Alvarado

## Abstract

Regeneration of body parts requires the replacement of multiple cell types. To dissect this complex process, we utilized planarian flatworms that are capable of regenerating any tissue after amputation. An RNAi screen for genes involved in regeneration of the pharynx identified a novel gene, Pharynx regeneration defective-1 (PHRED-1) as essential for normal pharynx regeneration. PHRED-1 is a predicted transmembrane protein containing EGF, Laminin G, and WD40 domains, is expressed in muscle, and has predicted homologs restricted to other lophotrochozoan species. Knockdown of PHRED-1 causes abnormal regeneration of muscle fibers in both the pharynx and body wall muscle. In addition to defects in muscle regeneration, knockdown of PHRED-1 or the bHLH transcription factor MyoD also causes defects in muscle and intestinal regeneration. Together, our data demonstrate that muscle plays a key role in restoring the structural integrity of closely associated organs, and in planarians it may form a scaffold that facilitates normal intestinal branching.

**Graphical Abstract:** 

**Highlights:** -PHRED-1 is a predicted transmembrane protein that contains Laminin G, EGF and WD40 domains

-PHRED-1 is required for normal muscle patterning during regeneration

-*phred-1* is expressed in muscle cells

-Muscle forms an essential scaffold for regeneration

## Introduction

Regeneration of complex organs requires the restoration of many cell types. For example, the regenerating salamander limb must regrow new osteoblasts, chondrocytes, muscle, neurons and epithelial cells simultaneously in order to allow complex tissues to regain form and function (Tanaka, 2016). Impairment of any one of these cell types will likely compromise regeneration of the whole organ. One of the challenges that regeneration biologists face is to understand how the sequential restoration of distinct tissues contributes to regeneration, and how different tissues interact during this process to coordinate replacement of complex structures.

Planarian flatworms are a powerful system for dissecting the molecular mechanisms of regeneration, due to their ability to regenerate any tissue rapidly after amputation (Reddien and Sánchez Alvarado, 2004) and the ease of testing gene function by RNA interference (Reddien et al., 2005; Rouhana et al., 2013). Regeneration is fueled by actively dividing stem cells called neoblasts (Rink, 2013). As stem cells divide, they differentiate into all the organs comprising the animal, including a central nervous system, an intestine, a pharynx for feeding, an excretory system, muscle, and epithelial cells (Roberts-Galbraith and Newmark, 2015). Although the instructive cues that direct differentiation into particular cell types are not known, recent studies have implicated muscle in the global patterning that occurs during whole-body regeneration (Lander and Petersen, 2016; Scimone et al., 2016; Witchley et al., 2013). Planarian muscle cells produce secreted molecules belonging to the FGF-receptor-like pathway and the Wnt pathway at distinct positions along the animal’s anterior-posterior axis. Together, these pathways are thought to create axial coordinates that pattern the anterior-posterior axis.

The planarian body is surrounded by a dense network of muscle fibers that provide structural support to the animal. This muscle network consists of distinctly patterned layers of fibers, including circular fibers that wrap around the body, longitudinal fibers that extend from anterior to posterior, and layers of diagonal fibers (Cebrià, 2016). The dorsal and ventral facets of the animal are connected by muscle fibers spanning the mesenchyme. Morphologically, although planarian muscle cells lack striations (Hori, 1983), they possess the molecular characteristics of striated muscle including thick filaments and striated muscle-specific myosin heavy chain (Kobayashi et al., 1998). Two conserved myosins have been described in planarians: myosin heavy chain-A (*mhcA*), expressed strongly in the pharynx, and myosin heavy chain-B (*mhcB*), expressed strongly in body wall muscle but not in the pharynx (Kobayashi et al., 1998; Orii et al., 2002). During regeneration, the muscle network extends over the blastema, initially appearing disorganized but gradually regaining its regularity and structure (Cebrià et al., 1997). The molecular cues required for patterning the restoration of muscle remain unclear.

We utilized an organ-specific regeneration assay, where we selectively removed the pharynx, and observed its regeneration, to identify genes that perturbed pharynx regeneration. Here we identify a previously unidentified gene to be required for normal muscle fiber formation both in the pharynx and in body wall muscle and name it *phred-1*, for *Ph*arynx *re*generation *d*efective-1. PHRED-1 is predicted to encode a transmembrane protein containing Laminin G, EGF, and WD40 domains. Among homologs in other animals, this particular domain arrangement occurs only in other lophotrochozoans. Inhibition of PHRED-1 resulted in defects in intestinal patterning, which we also observed after RNAi knockdown of other muscle effectors. Therefore, our findings have uncovered a novel role for muscle in planarian regeneration, and have implicated a previously undescribed protein in this process.

## Materials and Methods

### Animal husbandry

Asexual animals from the *Schmitea mediterranea* clonal line CIW4 were maintained at 20°C in Montjuïc salts (Newmark and Sánchez Alvarado, 2000). Animals were starved for ≥7 days prior to experiments. Planarians were irradiated on a J.L. Shepherd and Associates model 30, 6000 Ci Cs
^137^ instrument at approximately 5.90 Gy/min (17 min). Chemical amputations were performed as described in (Adler et al., 2014).

### RNAi and molecular biology

Initial RNAi was performed by bacterial administration as previously described (Reddien et al., 2005). Animals were fed three times over the course of 6 days, followed by chemical amputation and a feeding assay as described in (Adler et al., 2014). Additional RNAi experiments were done by injection of in vitro synthesized dsRNA, using MEGAscript T7 Transcription kit (AM1334, ThermoFisher). Injections were performed 3 times over the course of 6 days, followed by chemical or surgical amputation the next day. The *S. mediterranea* homolog of MHC-A was identified by BLAST and corresponds to the Planmine transcript dd_Smed_v6_579_0_2 (amplified with primers 5’-CGAAGTCCGAGAACATGCTCA-3’ and 5’-CAGGTGCTAATGTTCTTGCAG-3’). For MHC-A and MyoD RNAi experiments, dsRNA was synthesized as in (Rouhana et al., 2013) and fed to animals 6 times, 3 days apart, prior to amputation.

We used the original *phred-1* RNAi clone (NBE.8.11E, GenBank Accession AY967703) as a template for 5’ RACE, and confirmed an 8kb transcript corresponding to the Planmine transcript dd_Smed_v6_4789_0_1 (Brandl et al., 2016). The NCBI Conserved Domain Architecture Tool also identified several lophotrochozoan homologs with similar domain structure. Reciprocal BLAST with all homologs from other organisms confirmed that PlanMine dd_Smed_v6_4789_0_1 was the top hit.

For qRT-PCR experiments, total RNA was isolated from animals 5 days after the final injection of either unc-22, C-terminal, or FL-phred dsRNA. RNA was extracted by dissolving animals in Trizol (life Technologies), homogenizing them with an IKA homogenizer, and isopropanol precipitation. Superscript III (Life Technologies) was used to synthesize cDNA. PCR mixes were made with 2X SYBR mix (Applied Biosystems), run on an Applied Biosystems 7900HT, and results quantified using Ct methods. Primers used: phred-2_F ACGTGCCAGAAATTCTTTCC; phred-2_R CCCCAACATAAATGTGTCCA; phred-4_F TACATTGGGTGCCGGTTTAT; phred-4_R CCCCAACATAAATGTGTCCA; cyclophilin_F TTATTTGGCGATCTTGCTCC; cyclophilin_R TTTAAAACGTCCCCCATCTG

### Pharynx extraction and antibody staining

Following chemical amputation, pharynges were rinsed in planaria water and then fixed for 30 minutes in 4% paraformaldehyde. After rinsing they were incubated in block containing 0.5% horse serum diluted in PBSTx (PBS + 0.3% Triton-X-100). Primary antibody incubations were performed for 1-2 hours, or overnight, using these antibodies diluted in block: Acetyl-α-Tubulin rabbit monoclonal antibody (D20G3) #5335 (Cell Signaling Technology); phalloidin-Alexa-594 (ThermoFisher). Signals were developed using fluorescently-conjugated secondary antibodies from ThermoFisher Scientific. DAPI (Roche) was applied at 1:5000 dilution in PBSTx. Animals were imaged on an LSM 5LIVE or an LSM 510.

To quantify muscle fiber thickness, we drew straight lines perpendicularly across longitudinal muscle fibers in either the proximal or distal regions of confocal images of isolated pharynges. We then used the Plot Profile tool to obtain a graph of pixel intensity. Lines drawn at the base of the peaks measured the width of the peaks (corresponding to individual muscle fibers).

### In *situ* hybridization and immunohistochemistry

Colorimetric *in situ* hybridizations were performed as described in (Pearson et al., 2009) and fluorescent *in situ* hybridizations as in (King and Newmark, 2013). Antibody staining followed in situ hybridizations. For hematoxylin/eosin sections and immunohistochemistry, animals were fixed, embedded and prepared for sectioning as described in (Adler et al., 2014). For PHRED-1 antibody staining, a 1:5000 dilution of primary (in PBSTx containing 1% BSA) was preabsorbed with ∼100 fixed *phred-1(RNAi)* animals prior to staining wild-type animals, and developed with a goat anti-rabbit HRP secondary antibody. H3P staining was conducted as previously described (Adler et al., 2014) and quantified using Fiji. Animals were imaged on either a Zeiss Lumar Stereoscope or a Leica M165 Stereoscope equipped with a DFC7000T camera. All images were processed using Fiji.

### Antibody production and Western blot

To generate the PHRED-1 rabbit polyclonal antibody, a 100aa peptide from the extracellular region (sequence: EISKLNNFDLSQESLVLGFGKKSNGIVVELDDFKFLLTKTSKDYQSDSFQSEERYDTKSSQQERTINFNGQNYLKYNFENRIIRPSENEELDLQFKFSED) was synthesized and injected into rabbits by Strategic Diagnostics, Inc (Newark, DE). The serum was affinity-purified with the peptide antigen (by SDI) and used at a dilution of 1:5000 for immunohistochemistry. For whole-mount staining, the diluted antibody was preabsorbed with *phred-1(RNAi)* animals to remove background.

For western blots, lysis buffer containing 20mM Tris pH7.5, 100mM NaCl, 5mM EDTA, plus Halt protease inhibitors (ThermoFisher 78430) and PMSF was added (300μL to 10 animals) and animals were macerated with a plastic pestle in an eppendorf tube. Lysates were spun at max speed for 10 minutes, then supernatants were removed and quantified by Bradford assays.

Samples were run on 7.5% acrylamide gels and transferred onto PVDF in transfer buffer. Blots were blocked in 5% nonfat dry milk powder (all washes in TBST), then incubated with PHRED-1 antibody (1:1000), and goat anti-rabbit HRP (Zymed, discontinued antibody, 1:5000 dilution).

Signal was developed using Pierce ECL reagent on film and quantified using gel analysis tools in Fiji.

## Results

### Knockdown of phred-1 causes defects in pharynx muscle regeneration

Regeneration of the pharynx requires activation of stem cells and the production of diverse cell types (Roberts-Galbraith and Newmark, 2015). To identify genes necessary for organ regeneration, we used a strategy described in (Adler et al., 2014) and screened existing libraries of planarian cDNAs (Reddien et al., 2005).. Ten days after amputation, animals have completely regenerated the pharynx, and have regained the ability to readily ingest food (Figure 1A). We found that RNAi of one clone, NBE.8.11E, caused a significant reduction in feeding ability as compared to controls (Figure 1B). Closer examination of pharynx morphology by *in situ* hybridization with the pharynx-specific marker *laminin* revealed that after RNAi of NBE.8.11E, the pharynx failed to develop as well as controls, and exhibited a stunted, diamond-shaped morphology (Figure 1C).

**Figure 1.**
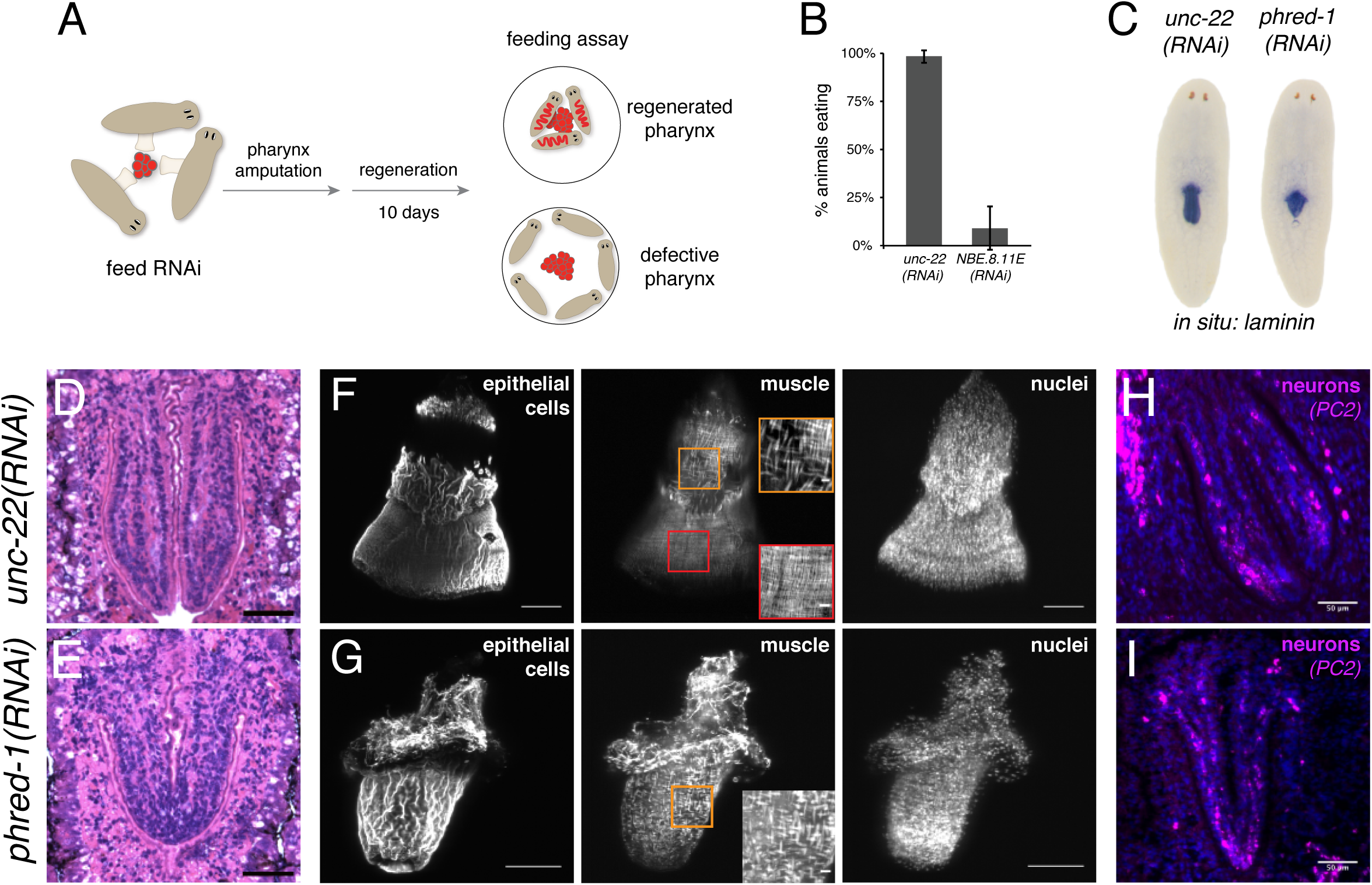
Pharynx regeneration requires PHRED-1. (A) Schematic of assay used to identify defective pharyngeal regeneration. (B) Percentage of animals able to ingest food 10 days after pharynx removal. (C) Whole-mount *in situ* hybridization for the pharynx marker laminin. (D-E) Paraffin sections from *unc-22(RNAi)* (D) or *phred-1(RNAi)* animals (E) stained with hematoxylin/eosin. (F-G) Isolated pharynges from control unc-22(RNAi) animals (F) or phred-1(RNAi) animals (G) stained with acetylated tubulin (ciliated epithelial cells), Tmus13 (muscle), and DAPI (nuclei). Insets show boxed regions from proximal (orange) and distal (red) regions. (H-I) Paraffin sections from *unc-22(RNAi)* (H) or *phred-1(RNAi)* animals (I) stained for the neuronal marker *PC2*. Scale bar, (D-E) 50μm, (F-G) 100μm, (H-I) 50μm.

The pharynx is a complex organ that consists of multiple cell types, including an extensive neural network (Cebrià, 2008; Okamoto et al., 2005), a coating of ciliated epithelial cells, and muscle. It is assumed that regeneration of a complete pharynx requires restoration of all of these cell types, to support normal feeding and chemotactic responses. Histological staining confirmed the constriction of the distal portion of the pharynx in *phred-1*(*RNAi*) animals as compared to controls (Figure 1D and 1E). To examine pharynx morphology at higher resolution, we extracted pharynges from animals with sodium azide, and then stained them with antibodies for muscle (Tmus13) and epithelial cells (acetylated tubulin). Pharynges from control animals had a bell shape and a smooth covering of ciliated epithelial cells, overlaying a fine network of muscle fibers (Figure 1F). Notably, a band at the proximal end of the pharynx fails to stain, suggesting that these cells are not ciliated. Epithelial staining of the pharynx highlighted two distinct morphological features: in the proximal region, the epithelium appeared ridged, while in the distal portion, the epithelial cells formed a smoother layer. Underlying these two regions, the muscle fibers also displayed differences: in the proximal region, muscle fibers appeared thicker than those in the distal region (4/4 animals, Figure 1F). Pharynges derived from *phred-1*(*RNAi*) animals were constricted at the distal end, and failed to form the distal region, exhibiting only thickened muscle fibers (6/6 animals, Figure 1G). To confirm that the distal pharynx fails to form, we examined two genes that are normally present in this region: the FGF receptor-like protein *ndl-4* and the secreted Frizzled-related protein *Sfrp-1*. In agreement with this defect in distal pharynx morphology, *ndl-4* and *Sfrp-1* exhibited altered expression in *phred-1(RNAi)* animals (Supplemental Figure 1). Despite this aberrant epithelial and muscle structure, the approximate number and distribution of neurons (as visualized with the neuronal marker *PC2*) was maintained in *phred-1*(*RNAi*) animals (Figure 1H and 1I; 34 vs 36.67 *PC2*^*+*^ cells in control vs. *phred-1(RNAi)* pharynges, n=3). Therefore, the morphological defects caused by knockdown of *phred-1* appeared to be restricted to the muscle and epithelial cells.

Pharyngeal muscle consists of three distinct types of fibers: longitudinal fibers running from the proximal to distal end, circular fibers that circumvent the pharynx, and radial fibers that connect the inner and outer lumens of the cylinder (Macrae, 1963) (Figure 2A, 2D, and 2G). Staining whole, extracted pharynges with the muscle monoclonal antibody Tmus13 (Bueno et al., 1997) revealed a network of fine, orthogonal fibers covering the surface of the pharynx in control animals (Figure 2B). By contrast, *phred-1*(*RNAi*) animals had thickened circular and longitudinal fibers (Figure 2C). Radial fibers connecting the inner and outer sheaths of the pharynx measured approximately 1-2μm in diameter (Figure 2E and 2H). In *phred-1*(*RNAi*) animals, these radial fibers were significantly thickened and failed to connect the inner and outer sheaths effectively (Figure 2F and 2I). Furthermore, quantification of longitudinal fibers showed that in control animals, muscle fibers are thicker in the proximal region than the distal region (Figure 2J). In *phred-1(RNAi)* animals, this difference is not evident (Figure 2J), confirming that the distal pharyngeal region fails to form. Based on the observed defects, we hypothesized that thickened muscle fibers and an improperly formed distal pharynx in *phred-1(RNAi)* animals resulted in an ineffective pharynx that cannot ingest food.

**Figure 2.**
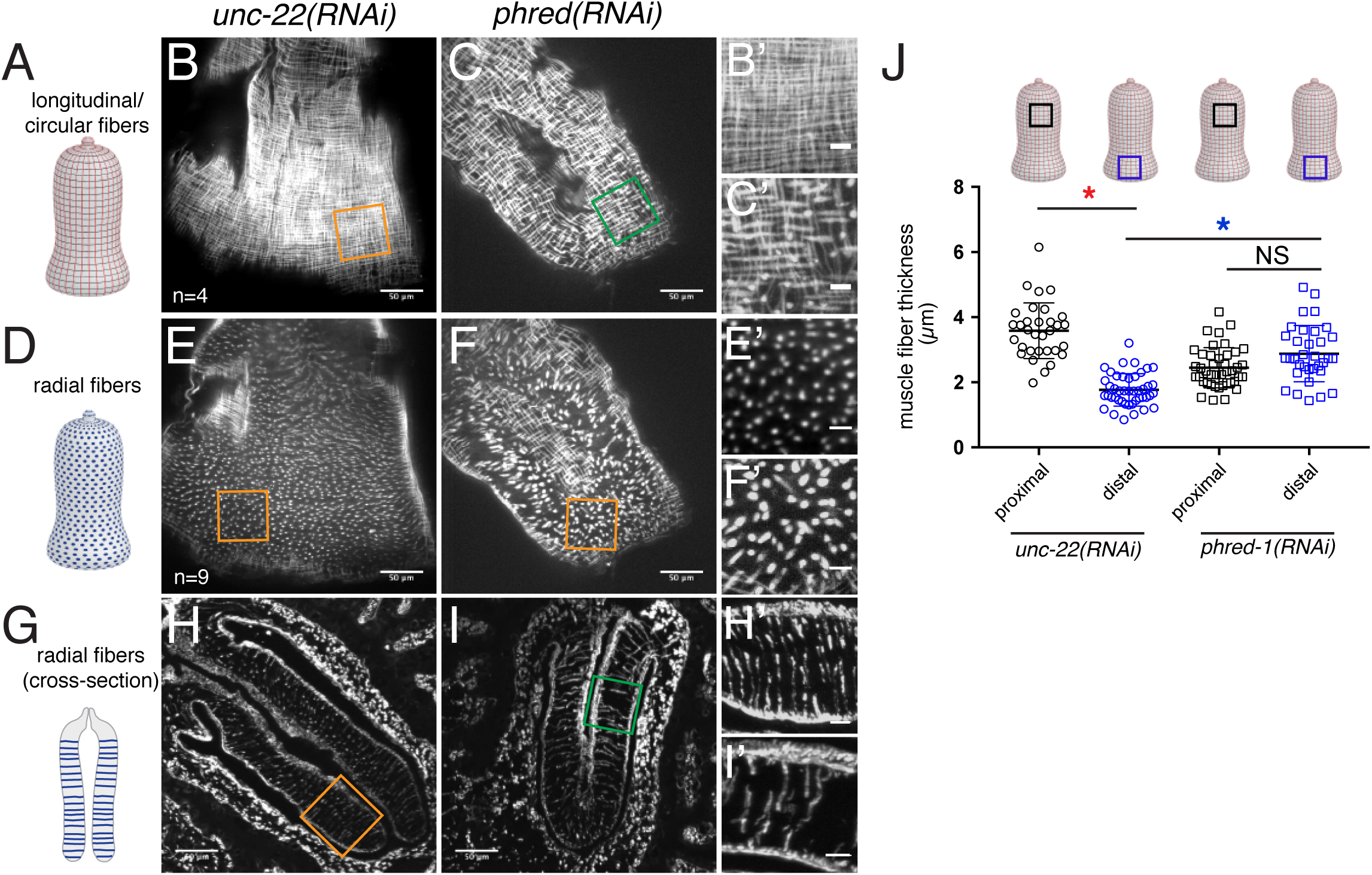
PHRED-1 is required for normal muscle fiber formation in the pharynx. (A-C) Longitudinal and circular fibers of the pharynx: schematic (A), isolated pharynges from *unc-22(RNAi)* (B) and *phred-1(RNAi)* (C) animals. Zoomed regions (orange boxes) are shown in B’, C’, E’, F’, H’, and I’. (D-F) Radial muscle fibers connecting inner and outer edges of the pharynx: schematic (D), isolated pharynges from *unc-22(RNAi)* (E) and *phred-1(RNAi)* animals (F). (G-I) Radial fibers in cross-section: schematic (G), paraffin sections from *unc-22(RNAi)* (E) and *phred-1(RNAi)* animals (F) stained with Tmus13 antibody. All animals imaged by confocal 14 days after amputation. Scale bars, (B-C, E-F, H-I) 50μm; (B’-C’, E’-F’, H’-I’) 10μm. (J) Quantification of muscle fiber thickness in isolated pharynges. Red asterisk compares *unc-22(RNAi)* proximal and distal muscle fibers; blue asterisk compares distal muscle fibers in *unc-22(RNAi)* and *phred-1(RNAi)* animals. Asterisks represent p<0.0001, two-tailed Students’ t-test. NS, not significant.

### PHRED-1 is a large transmembrane protein expressed in muscle

Sequencing of the NBE.8.11E RNAi clone identified a 3kb gene containing a single laminin G domain, an EGF domain, a predicted transmembrane region, and three WD40 domains (Figure 3A), but no strong homology to other known proteins. Because the predicted protein appeared to span the membrane but lacked a signal sequence, we performed 5’-RACE to determine its full-length sequence, and verified predicted transcripts with recent *S. mediterranea* transcriptomes (Brandl et al., 2016). We identified the full-length gene as an 8kb transcript, which encodes a 2536 aa protein (Figure 3A). At its N-terminus, PHRED-1 contains a predicted signal sequence. On its extracellular side, PHRED-1 contains 6 Laminin G domains and 9 EGF domains. These domains are commonly found in cell adhesion proteins, but this particular combination occurs in proteins involved in cell-cell contact, including Crumbs, contactin, and neurexin. SMART (Letunic et al., 2015; Schultz et al., 1998) identified 3 intracellular WD40 domains, common protein-protein interaction domains involved in a wide variety of cellular processes (Xu and Min, 2011). BLAST with full-length PHRED-1 identified several direct homologs with similar domain structure (Figure 3A). However, the only animals with direct homologs containing Laminin G, EGF, and WD40 domains together belong to the lophotrochozoan clade (including the parasitic flatworm *Schistosoma mansoni*, the limpet *Lottia gigantea*, Octopus, and the polychaete *Capitella*) (shown in Supplemental Figure 2A).

**Figure 3.**
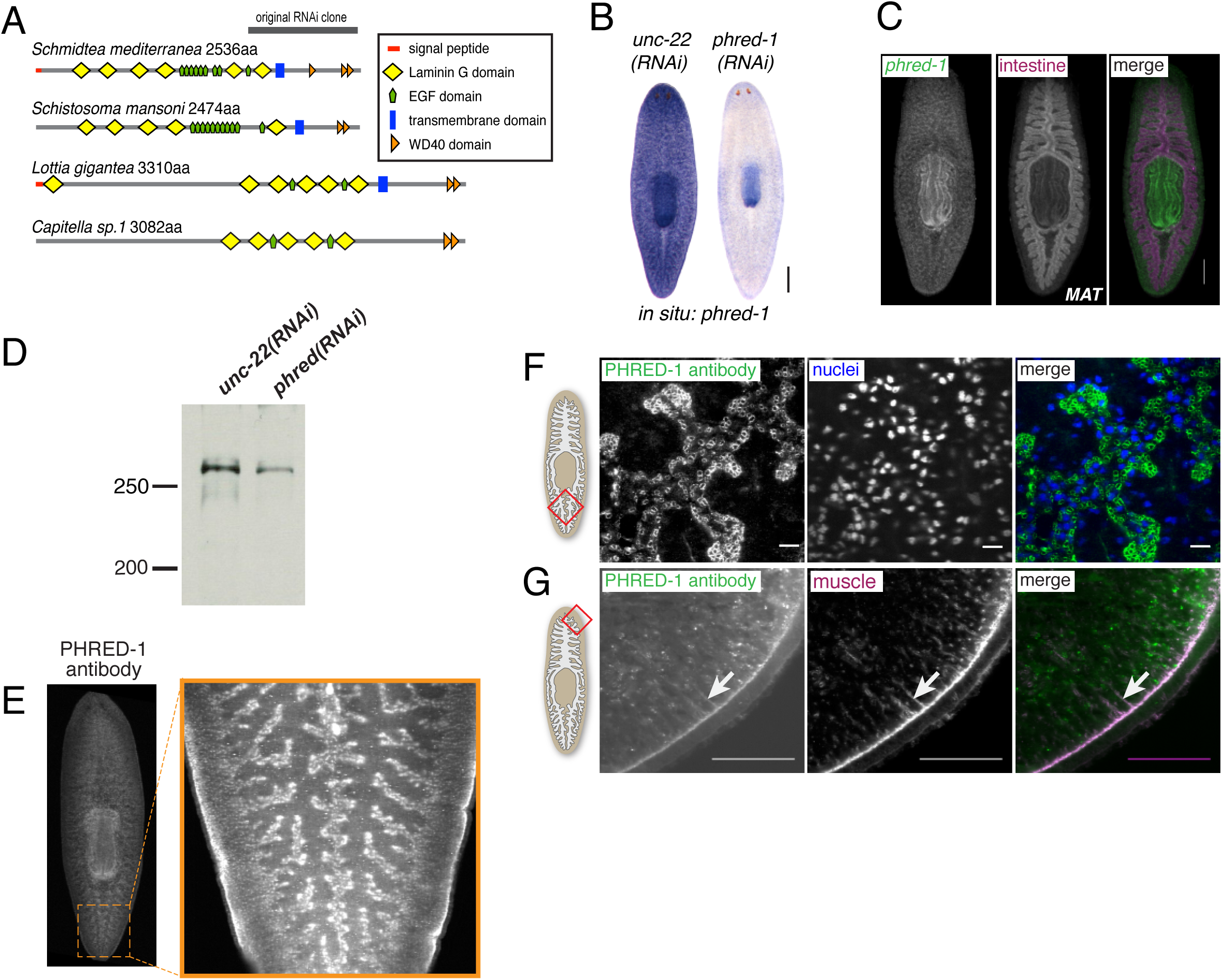
PHRED-1 is a novel transmembrane protein expressed in muscle cells. (A) Domain prediction of PHRED-1 and other lophotrochozoan homologs. (B) WISH of *phred-1* in control and *phred-1(RNAi)* animals. (C) Double fluorescent *in situ* hybridization for *phred-1* and *MAT* (intestinal marker). Western blot of lysates from control *unc-22(RNAi)* and *phred-1(RNAi)* animals. Predicted molecular weight of PHRED-1 is 298kD. (E) Whole-mount immunostaining with PHRED-1 antibody. Zoomed region shows staining outside of intestine. (F) Immunohistochemistry of tail region (red box on schematic). Paraffin sections stained with PHRED-1 antibody and DAPI, highlighting small circular structures, likely myofibers, between instestinal branches. (G) Paraffin sections of peripheral region (red box on schematic) stained with PHRED-1 and Tmus13 (muscle) antibodies. White arrows highlight radial fibers. Scale bar, (B-C): 250μm. (F-G) 50μm.

Alignments with MUSCLE highlighted that the C-termini of these family members is strongly conserved throughout two WD-40 domains, with significant identity in primary sequence (Supplemental Figure 2B). Outside of lophotrochozoans, this combination of Laminin, EGF, and WD40 domains does not occur in the same protein. Among the available lophotrochozoan homologs, none have been studied at a functional or descriptive level.

*In situ* hybridization of *phred-1* indicated strong expression in both the pharynx and the rest of the animal (Figure 3B). *phred-1* RNAi knockdown caused a strong reduction in transcript levels, visualized by *in situ* hybridization and by qRT-PCR (Supplemental Figure 3). Analysis with other anatomical markers highlighted that *phred-1* is expressed outside of the intestine (Figure 3C), in a pattern reminiscent of *myosin heavy chain A* (*DjMHC-A*) expression (Cebrià, 2016; Kobayashi et al., 1998), a myosin specifically expressed in planarian body wall musculature. We raised a polyclonal antibody against the extracellular portion of the protein, which has a predicted molecular weight of 298 kD. We verified the specificity of this antibody by Western blot and found that a single, large protein with molecular weight >270kD had reduced signal in of *phred-1(RNAi)* animals as compared to controls (36% knockdown relative to nonspecific bands, Figure 3D). Using this antibody, we confirmed that PHRED-1 protein shows a similar distribution pattern to its transcript, enriched in the pharynx and outlining the intestine (Figure 3E). In our immunohistochemistry, we noticed that the PHRED-1 antibody labeled small, round structures nestled between the intestinal branches (Figure 3F, region shown clearly in Figure 3E). Because these structures were smaller than nuclei, we hypothesized that these were muscle fibers, and confirmed that PHRED-1 colocalized with muscle markers such as the monoclonal antibody Tmus13 (Bueno et al., 1997) (Figure 3G). In addition, we searched a database of planarian single-cell RNA-sequencing data with *phred-1* (Wurtzel et al., 2015), and found that *phred-1* clearly localized among other muscle-expressed genes (Supplemental Figure 4). Together, these results suggest that PHRED-1 is a protein expressed on the surface of muscle cells. Given its conserved domain structure, we hypothesize that PHRED-1 may play a role in cell-cell interactions.

**Figure 4.**
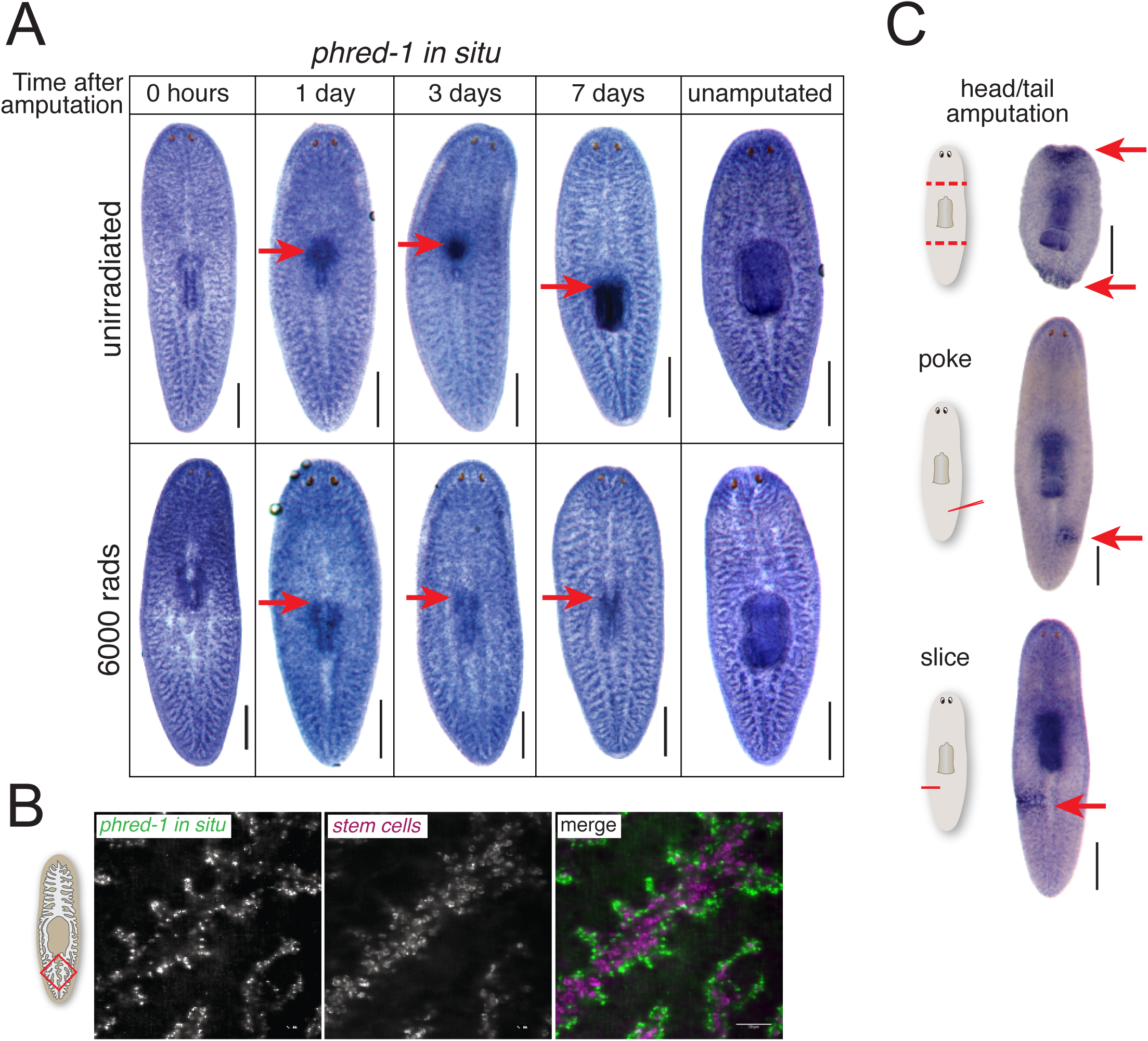
Wound-induced *phred-1* expression in muscle cells. (A) *phred-1 in situ* hybridization in animals at specified times after pharynx removal in either (A) unirradiated animals (top) or animals exposed to 6,000 rads of gamma-irradiation (bottom). (B) Fluorescent *in situ* hybridization showing *phred-1* and *piwi-1* expression in the tail region (red box on schematic). (C) *phred-1 in situ* hybridization in animals injured as indicated and fixed 12 hours later. Scale bars, (A) 500μm. (B) 50μm.

Muscle cells drive contraction of wounds (Chandebois, 1979), an essential step in initiation of healing and regeneration. Although *phred-1(RNAi)* animals had no obvious defects in wound healing, we considered the possibility that injury might alter *phred-1* transcription during pharynx regeneration. We noticed that the *phred-1* transcript increases strongly in the vicinity of the regenerating pharynx within 24 hours of amputation (Figure 4A), and is sustained in the newly produced organ. We previously showed that a burst of stem cell proliferation occurs as the pharynx initiates regeneration (Adler et al., 2014), and stem cell progeny begin expressing pharynx-specific markers 1-2 days after amputation. To test whether stem cell progeny were responsible for expressing *phred-1*, we blocked regeneration by eliminating stem cells with lethal doses of radiation (Eisenhoffer et al., 2008). Under these conditions, we observed no local increase in *phred-1* expression (Figure 4B), indicating that without formation of new tissue, *phred-1* is not expressed, but its expression in muscle cells in the rest of the animal remains unaltered.

Planarian stem cells reside in the mesenchyme which outlines the intestine, in a pattern similar to the distribution of *phred-1*. To determine whether *phred-1* was expressed in stem cells, we tested its colocalization with the stem cell marker *Smedwi-1*. We found almost no overlap between these two transcripts (Figure 4B). Supporting the absence of colocalization with stem cells, the expression pattern of *phred-1* was unchanged after lethal doses of radiation (Figure 4A). Additionally, searching a database of planarian single-cell RNA-sequencing data failed to strongly place *phred-1* among neoblasts (Wurtzel et al., 2014; Supplemental Figure 4B). We conclude that *phred-1* is expressed in both planarian body wall and pharyngeal muscles.

To test whether *phred-1* expression was induced by other types of wounds, we amputated animals in several different ways. Twelve hours after head and tail amputation, we noticed a rapid increase in *phred-1* expression (Figure 4C). Even small incisions or a needle poke – injuries that do not stimulate local increases in neoblast activity (Baguñà, 1976; Wenemoser and Reddien, 2010) – activated *phred-1* expression at this time point (Figure 4C). Therefore, body wall muscle cells respond to injury by increasing expression of *phred-1*.

### Muscle forms an important scaffold for whole-body regeneration

Planarian regeneration requires concurrent regrowth and patterning of multiple organs. For example, head fragments must regenerate a new pharynx at the posterior end while simultaneously extending two intestinal branches around it, toward the tail (Forsthoefel et al., 2011). Because *phred-1* is broadly expressed throughout the body, we tested whether PHRED-1 is required for whole-body regeneration in addition to pharynx regeneration. We found that regeneration of posterior regions was compromised in head fragments. *phred-1(RNAi)* animals failed to form posterior blastemas (2/13 animals as compared to 6/6 in control animals) (Figure 5A). Surprisingly, despite this strong defect in posterior blastema formation, tail fragments regenerated heads normally (8/9 as compared to 8/8 in control animals, not pictured). Similarly, trunk fragments also appeared to regenerate normally. However, after feeding, the anterior intestine filled excessively with colored food, becoming distended and overfilled (Figure 5B).

**Figure 5.**
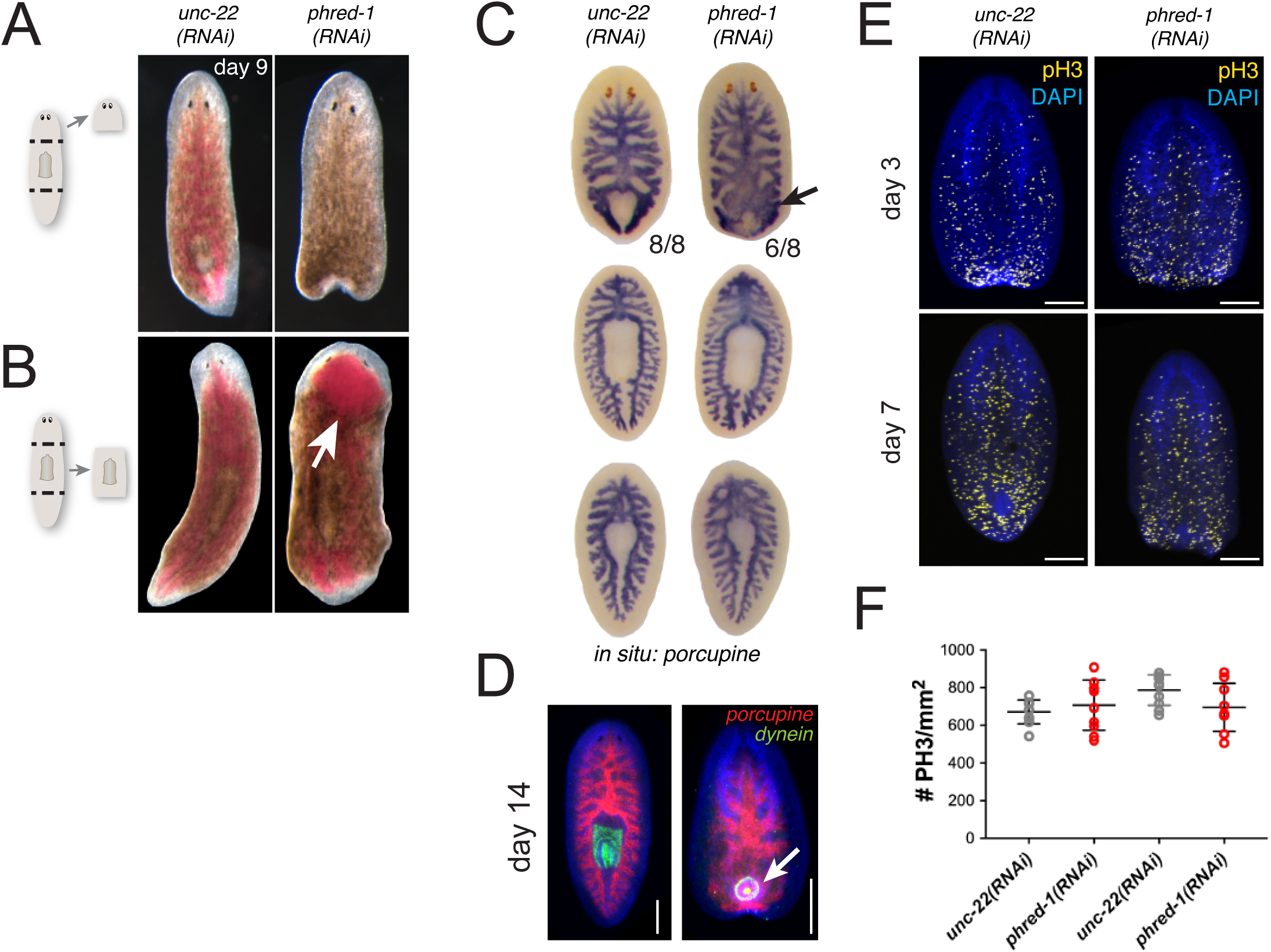
PHRED-1 is required for normal intestinal morphogenesis during regeneration. (A) Head fragments 9 days after amputation. *phred-1(RNAi)* heads fail to regenerate posterior ends. (B) Trunk fragments 9 days after amputation. After feeding, the anterior end of the intestine overfills with food (arrow). (C) *In situ* hybridization with the intestinal marker porcupine in fragments 7 days after amputation. Arrow highlights aberrant branching in *phred-1(RNAi)* animals. (D) Fluorescent *in situ* hybridization with porcupine (red) and dynein heavy chain (green), showing defective pharynx positioning in *phred-1(RNAi)* animals. (E) Phosphohistone-H3 staining in head fragments 3 and 7 days after amputation. (F) Quantification of phospho-histone-H3 staining in head and tail fragments 7 days after amputation. Scale bars (D-E), 200μm.

Together with the prior observation that PHRED-1 may regulate muscle, the defect in intestinal filling suggests that the body wall muscle surrounding the intestine may limit intestinal distortion after feeding. Examination of intestinal morphology revealed that head fragments exhibited thickened anterior branches and fusion of secondary branches in the vicinity of the pharynx (Figure 5C and 5D). *phred-1* is expressed in the body wall musculature surrounding the intestine, suggesting the possibility that planarian body wall muscle could form a scaffold that envelops the intestine. The tight association between the muscle and the intestine may facilitate intestinal branching and provide structural support for the intestine after food ingestion.

Because regeneration depends on stem cell proliferation (Newmark and Sánchez Alvarado, 2000; Rink, 2013), we first examined the extent of proliferation during regeneration using the mitotic marker phosphohistone-H3. No significant differences were observed when we quantified the number of PH3-positive nuclei in control and *phred-1(RNAi)* animals (Figure 5E and 5F). This result suggests that impaired stem cell proliferation is not responsible for the defects observed in *phred-1(RNAi)* animals. Instead, PHRED-1 is likely to control either cell-cell interactions or muscle cell morphology that facilitate regeneration.

To determine whether PHRED-1 is involved in muscle regeneration, we monitored muscle morphology in regenerating blastemas of head fragments. We directly visualized muscle cells with the monoclonal antibodies Tmus13 or 6G10 (Bueno et al., 1997; Ross et al., 2015). Control animals had a dense, continuous network of muscle fibers in the blastema, extending to the distal edge of the animal (Figure 6A). By contrast, *phred-1(RNAi)* animals displayed disarrayed muscle fibers, with the blastema appearing frayed and disintegrating (Figure 6A). To confirm that this defect was due to a muscle deficiency, we knocked down the bHLH transcription factor MyoD, a key regulator of muscle cell differentiation (Tapscott, 2005). In planarians, *myoD* knockdown causes defects in brain regeneration and blastema patterning (Cowles et al., 2013; Reddien et al., 2005). Knockdown of *myoD* caused qualitatively similar defects to *phred-1*(*RNAi*) animals, with a disorganized muscle fiber network in the blastemas of regenerating head fragments (Figure 6A). We observed similar defects in tail fragments (data not shown). These data support the function of PHRED-1 in driving muscle cell regeneration.

**Figure 6.**
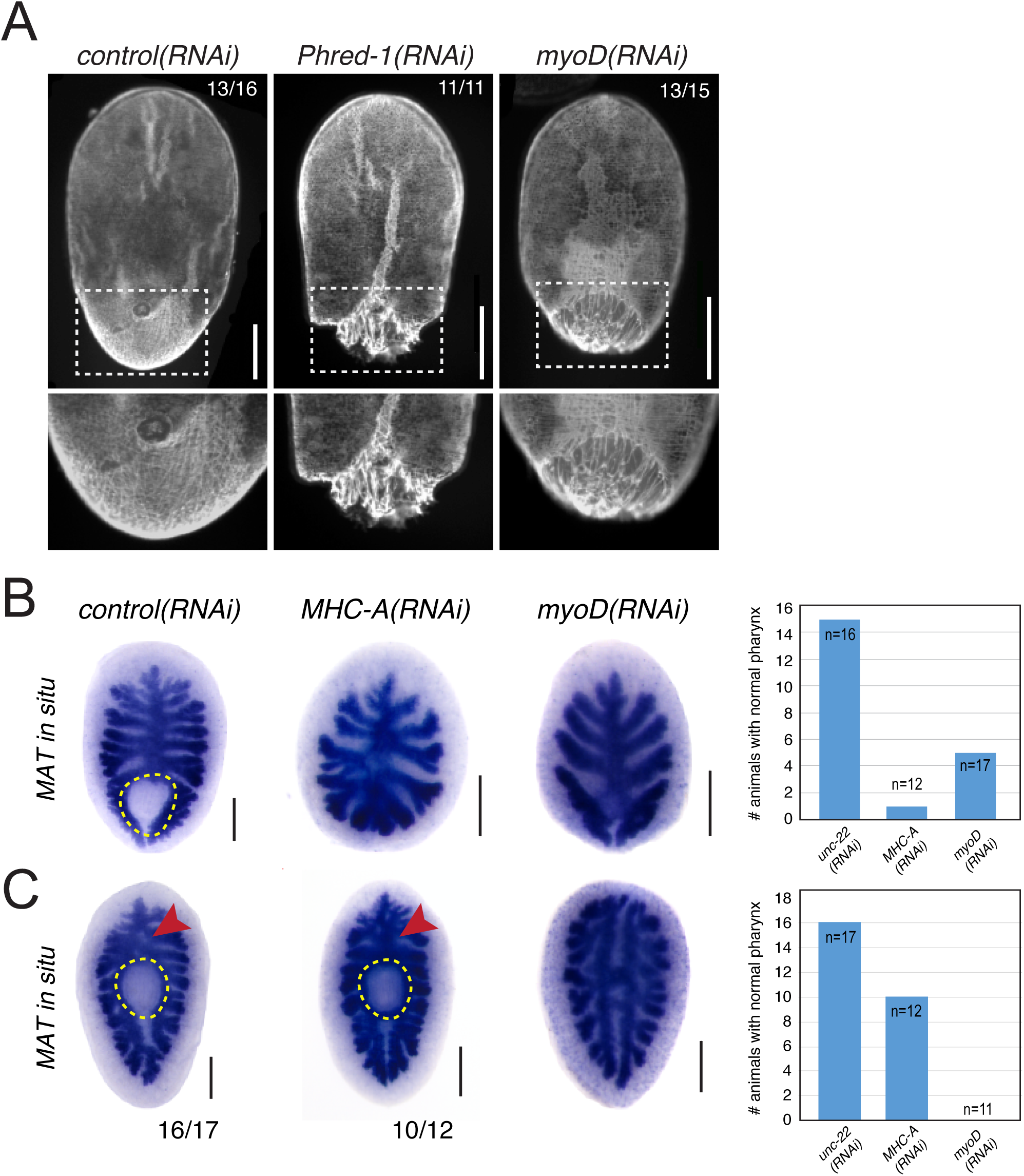
Muscle forms an essential scaffold for regeneration. (A) Head fragments from RNAi animals stained with 6G10 (muscle) antibody. Numbers indicate animals with the morphology shown from one representative experiment. Boxes highlight region below. (B-C) *In situ* hybridizations with the intestinal marker *MAT*. Head fragments (B) and tail fragments (C) 7 days after amputation. Pharynx region is highlighted by dashed yellow line. Red arrowheads highlight the primary intestinal branch. Quantification of fragments with a regenerated pharynx are shown on right, with population sizes indicated. Scale bars, 250μm.

Because of the intestinal patterning defects we observed after *phred-1* knockdown (Figure 5), and due to the fact that body wall muscle surrounds the entire intestine, we hypothesized that body wall muscle might play a structural role in regeneration. We tested this hypothesis by examining intestinal morphology in animals knocked down for either the myosin heavy chain gene *mhcA*, which is strongly expressed in the pharynx and body wall muscle like *phred-1* (Kobayashi et al., 1998), or the myogenic transcription factor *myoD*. Although knockdown of these genes was incomplete (Supplemental Figure 5), both *mhcA(RNAi)* and *myoD(RNAi)* animals exhibited defective intestinal regeneration in head and tail fragments (Figure 6B and 6C). In addition, both *mhcA(RNAi)* and *myoD(RNAi)* animals failed to form a pharynx. These results demonstrate that muscle is an essential component of the regeneration blastema that facilitates normal patterning of underlying organs.

The shared phenotypes of *mhcA*, *phred-1*, and *myoD* raised the possibility that muscle genes generally respond to wounding, similar to *phred-1*. Twelve hours after wounding, we observed that *mhcA* transcript levels increase at the wound site, but *myoD* does not respond at all (Supplemental Figure 6A-B). Furthermore, collagen, a known marker of muscle cells in planarians (Witchley et al., 2013), does not increase at the wound site (Supplemental Figure 6C). Together, these results suggest that muscle cells can exhibit a variety of responses to wounding.

### Muscle cells are important for anterior-posterior patterning

In *phred-1(RNAi)* animals, head fragments form a small pharynx as compared to controls (Figure 3F), suggesting a failure in the repatterning process that occurs during regeneration. Recent studies have shown that anterior-posterior patterning in planarians is controlled in part by a network of signaling molecules expressed in distinct positions from head to tail (Scimone et al., 2016; Witchley et al., 2013). To test whether the observed *phred-1(RNAi)* defects were due to alterations in the muscle-expressed patterning genes, we evaluated the expression of these genes in animals with defective musculature. The gene encoding a secreted Wnt known as *wnt11-5 (*also called *wntP-2)* is normally expressed in the posterior region of animals, spanning the region between the anterior of the pharynx and the tip of the tail (Gurley et al., 2010; Petersen and Reddien, 2009). Amputation causes a disruption of the normal distribution of *wnt11-5* expression, resulting in abnormally low levels in head fragments and high levels in tail fragments. As the anterior-posterior axis resets during regeneration, *wnt11-5* expression gradually becomes excluded from the anterior of regenerating fragments, concentrating in the posterior. In regenerating *phred-1(RNAi)* fragments, the domain of *wnt11-5* is slightly increased over controls (Figure 7A and 7C). Strikingly, in *myoD(RNAi)* tail fragments, which have strong defects in brain regeneration and fail to form a pharynx (Figure 6C), *wnt11-5* expression extends all the way to the anterior tip (Figure 7B and 7D), occupying the entire regenerating fragment and not becoming excluded from the anterior region. The failure to reset *wnt11-5* expression occurs throughout the fragment, in both the newly regenerated muscle cells at the anterior, as well as in the pre-existing muscle in the remainder of the fragment. These results suggest that inhibition of muscle can perturb the normal anterior-posterior expression of patterning genes, and may have direct consequences on regeneration of underlying organs like the brain.

**Figure 7.**
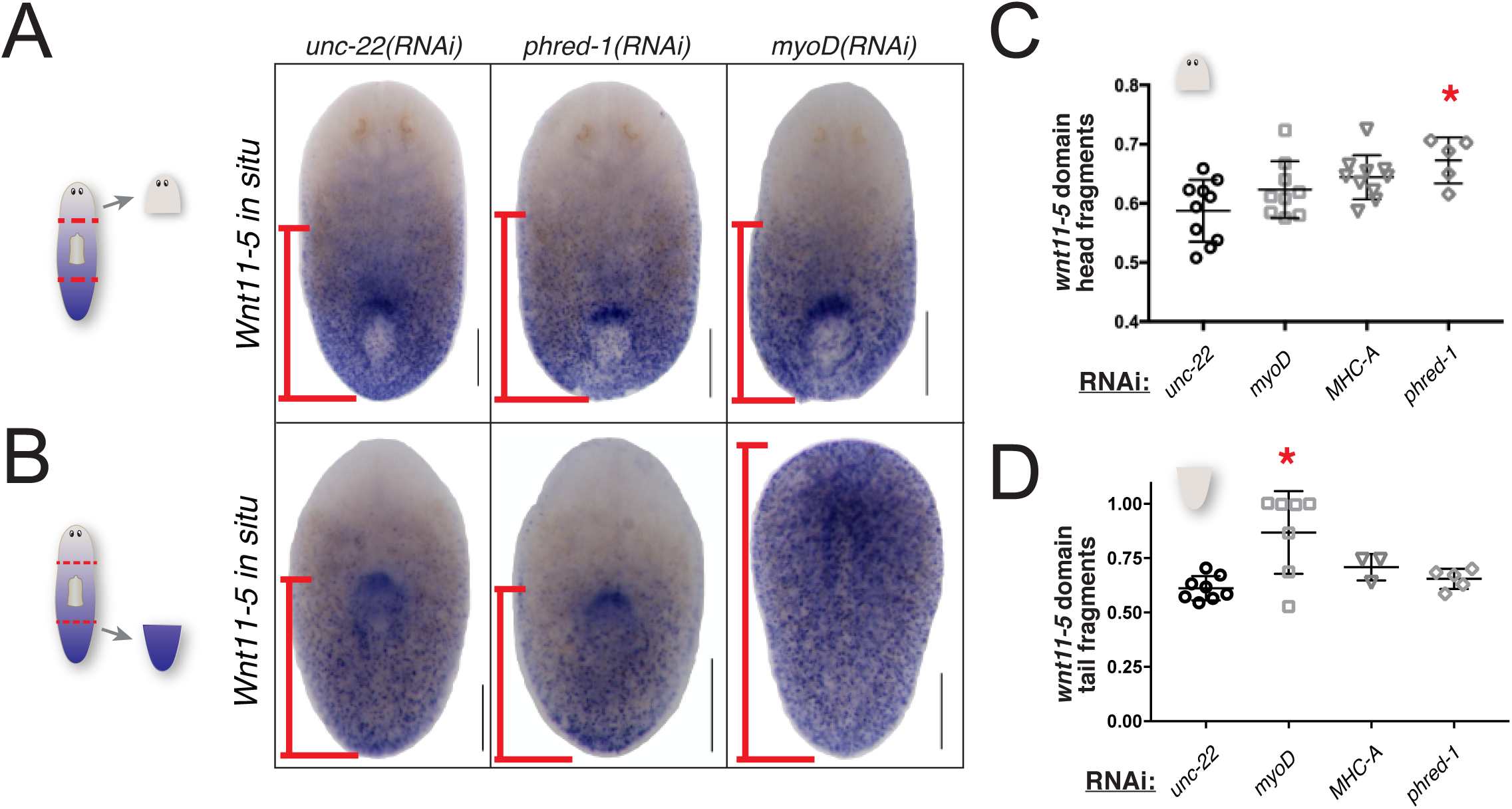
Muscle is required for expression of axial patterning genes. (A-B) *wnt11-5 in situ* hybridization in regenerating head fragments (A) and tail fragments (B), 7 days after amputation. Scale bars, 250μm. Red lines show domain of *wnt11-5* quantified in C and D. (C-D) Quantification of the domain occupied by *wnt11-5* expression, relative to the total length of the fragments. Head fragments (C) and tail fragments (D), 7 days after amputation. Asterisks mark significant differences as compared to *unc-22(RNAi)* controls (P<0.005, Student’s two-tailed t-test). Scale bars, 250μm.

## Discussion

Regeneration of complete organs or entire body parts requires the restoration of multiple tissue types. One of the main goals of regeneration biology is to understand the timing and relationships between cell types as they are replaced (Tanaka, 2016). Here, we have identified a lophotrochozoan-specific protein that is required for various aspects of regeneration in planarians. PHRED-1 is a predicted transmembrane protein expressed in muscle, which patterns normal muscle fibers during regeneration. We found that proper muscle fiber formation is essential for establishing normal morphology of the intestine during regeneration. We confirmed this new role for muscle by knocking down central regulators of myogenesis (the bHLH transcription factor MyoD) and function (myosin heavy chain A, MHC-A), both of which also caused intestinal regeneration and patterning defects similar to those observed in *phred-1(RNAi)* animals. Therefore, we have identified a novel role for muscle in planarian regeneration, as well as a new molecule involved in this process.

### Putative signaling roles for PHRED-1

The unique assortment of domains in a protein confers its distinctive interactions with extracellular proteins, intracellular proteins, and facilitates signal transduction (Basu et al., 2008). The extracellular region of PHRED-1 consists of several laminin G and EGF domains, which are commonly found in other proteins known to be important during development, such as Neurexin, Notch and Crumbs. However, the addition of WD40 domains in the intracellular region demarcates PHRED-1 and its homologs in other lophotrochozoans as a distinct protein family.

Over the course of evolution, protein domains have been observed to undergo shuffling (Jin et al., 2009), which allows them to interact with new partners, and potentially to acquire novel functions. How this divergence occurred in a lophotrochozoan ancestor is unclear. Importantly, the functional attributes due to the addition of these WD40 domains remain to be elucidated. In this study, we only investigated the function of PHRED-1 in adult animals, but it is equally possible that PHRED-1 acts during embryonic development to guide formation or function of muscle.

Recent characterization of muscle in the marine annelid *Platynereis dumerilii* and *Schistosoma mansoni* has highlighted that other lophotrochozoans possess muscle cells that cannot be clearly classified as either smooth or striated (Brunet et al., 2016; Sulbarán et al., 2015). These hybrid muscle cells may harbor a unique set of effectors facilitating their function. The evolutionary novelty of PHRED-1 could be due to the unique morphological and molecular characteristics of muscle cells within lophotrochozoans, and PHRED-1 may be one of the downstream effectors of this particular muscle cell type. Because PHRED-1’s closest homologs in non-lophotrochozoan animals are involved in cell-cell contact, it is possible that PHRED-1 may function in mediating contact between these unusual muscle cells.

*phred-1* knockdown does not appear to cause regeneration defects by impairing stem cell function, like a recently described matrix metalloproteinase (Dingwall and King, 2016). Instead, it appears to limit the thickness of myofibrils. Electron microscopy has shown that myofilaments within the pharynx can vary in width (Macrae, 1963). Knockdown of *phred-1* causes defects in muscle fiber morphology, specifically thickening and disrupting their patterning. RNAi-based screens in *C. elegans* have identified several molecules causing similar phenotypes (Meissner et al., 2009), but the functions of these genes are diverse. No obvious homologs of *phred-1* emerged in analyses of muscle formation in other animals. However, future work may identify other genes shared between ecdysozoans and lophotrochozoans in regulating muscle patterning. Because planarian muscle does not have a dedicated satellite cell population as vertebrates do (Cebrià, 2016), understanding the differences between muscle formation during development and muscle restoration after injury will be important avenues to pursue.

### Role of muscle in regeneration: wound healing, patterning, or structural scaffold?

Minutes after an amputation, muscle contractions reduce the size of the wound (Chandebois, 1979), followed by migration of epithelial cells over the injured tissue. Once the wound is closed, blastema formation begins and muscle fiber structure is restored, eventually becoming indistinguishable from the pre-existing tissues (Cebrià and Romero, 2001).

Knockdown of *phred-1*, *mhcA,* and *myoD* did not appear to affect wound closure, suggesting either that these proteins do not affect the acute response of muscle cells to injury, or that sufficient protein perdures in uninjured tissue to facilitate wound closure (see below). Many genes are known to robustly increase after wounding in planarians (Wenemoser et al., 2012), but the function of this transcriptional increase in wound healing remains elusive. The contractions that occur after injuries could also initiate transcriptional changes. The increase in *phred-1* expression that occurs hours after wounding suggests that muscle cells respond to injury by upregulating gene expression. In fact, a recent study showed that muscle cells have a robust transcriptional response after wounding, detected by single-cell RNA-sequencing (Wurtzel et al., 2015). The function of this transcriptional response has yet to be determined.

Expression analysis of planarian muscle genes reveals that the planarian muscle system is a complex tissue. Despite the fact that previous analysis of purified neoblasts identified *myoD* and *collagen* in the same cell, their expression patterns are very different in the animal (Cowles et al., 2013, Scimone et al., 2014b), suggesting that they may diverge during muscle cell differentiation. Similarly, our analysis of gene expression changes after wounding demonstrates that muscle cells do not respond uniformly to injuries. *phred-1* and *mhcA* increase expression while *myoD* and *collagen* do not. Whether these differences depend on unique functions of genes, or their expression within particular subsets of muscle cells is still unclear. Additionally, the incomplete knockdown observed for *phred-1*, *myoD* and *mhcA* precludes our ability to fully assess how muscle contributes to wound healing. The partial knockdown we observe may be a result of long-lived muscle cells, a resistance to RNAi knockdown, or a byproduct of the complexity of planarian muscle.

The most obvious defects caused by inhibition of muscle occur in the newly produced tissue, an observation consistent with prior myosin knockdowns (Sánchez Alvarado and Newmark, 1999). The thickened muscle fibers and frayed blastemas that we observed after knockdown of *phred-1*, *mhcA,* and *myoD* suggest that regeneration demands significant production of new muscle cells. The regeneration blastema consists entirely of newly produced cells, and muscle cells cannot re-establish their normal orthogonal network. The absence of defects in muscle organization in pre-existing tissue and uninjured animals suggests that muscle cells either do not undergo rapid turnover, or that intercalation of single muscle cells into an existing lattice does not require the same signaling pathways.

Recent evidence has suggested that planarian body wall muscle cells produce signaling molecules that govern anterior-posterior patterning during regeneration (Scimone et al., 2016; Witchley et al., 2013). These molecules, many of which are members of the Wnt and FGF signaling pathways, are expressed in distinct domains along the anterior/posterior axis. Amputation causes drastic changes in the normal distribution of these molecules, but as regeneration progresses, they re-establish their normal expression domains (Gurley et al., 2010; Petersen and Reddien, 2009). In uninjured animals, *wnt11-5* is restricted to the posterior of the animal, with strong signal at the pharynx. Amputation causes drastic alterations in the gradient of *wnt11-5* expression, and as the animal regenerates, *wnt11-5* gradually becomes excluded from the anterior region. Tail fragments from *myoD(RNAi)* animals fail to suppress *wnt11-5* expression in the anterior region of tail fragments, and also completely fail to regenerate a new pharynx. How pharynx regeneration may be controlled by these axial patterning genes remains unclear, although recent reports suggest the involvement of the Wnt co-receptor *Ptk7* and the FGF receptor-like molecule *ndl-3* (Lander and Petersen, 2016; Scimone et al., 2016). The impaired ability to reset expression of anterior-posterior signaling molecules may explain the strong regeneration defects observed in *myoD(RNAi)* animals (Cowles et al., 2013; Reddien et al., 2005). Additionally, MyoD could be important for establishing the anterior pole in tail fragments, which requires the Wnt inhibitor notum, the TALE transcription factors PREP and PBX, the transcription factor ZicA, and the Forkhead transcription factor FoxD (Vogg et al. 2014;

Petersen and Reddien 2011; Scimone et al. 2014a; Felix and Aboobaker 2010; Blassberg et al. 2013; Chen et al. 2013; Vásquez-Doorman and Petersen 2014).

The planarian intestine is the largest organ in the body. Just outside of the intestine are muscle fibers that connect the dorsal and ventral sides of the animal. How this complex network of integrated myofilaments is restored after injury, and how they may interact with the regenerating, branched intestine, is not known (Cebrià, 2016). The distortion of intestinal regeneration we observed after knockdown of *phred-1*, *mhc-A,* and *myoD* suggests that muscle may form a structural support for regeneration of other organs or cell types. Therefore, without an organized network of myofilaments, regeneration stalls. As planarians regenerate complex structures comprised of multiple tissue types, the patterning of one tissue (*e.g.*, the intestine) may rely on the sequential replacement of other cell types. The dependence of one tissue on another during regeneration also occurs in the pharynx, where formation of this muscular scaffold appears to be an essential foundation for subsequent assembly of its neuronal and secretory components. Furthermore, how muscle and intestinal branches interact during regeneration, and whether an interaction between muscle and intestinal branches contributes to its elaborate and dynamic branching patterns (Forsthoefel et al., 2011) will be interesting to elucidate.

## Conclusions

We identified a putative transmembrane protein that is important for myofilament formation in planarian muscle cells. PHRED-1 contains laminin G, EGF, and WD40 domains, and is present in all muscle cells. Our findings reveal that planarian musculature coordinates whole-body regeneration in two ways: first, by producing signaling molecules that pattern the regeneration of complex structures, and second, by providing necessary structural support to surrounding organs. Organ regeneration relies on the accurate replacement, organization, and functional integration of the specific cell types required for each unique structure. Our work demonstrates that restoration of musculature is an essential component of organ regeneration. Future work will determine whether analogous proteins have similar functions in the musculature of other animals.

## Acknowledgements

We would like to thank the Histology Core at the Stowers Institute for Medical Research for sectioning support, and Divya Shiroor for critical reading of the manuscript. This work was supported by NIH Developmental Biology Training Grant 5T32 HD07491 and NRSA F32GM084661 (C.E.A.) and NIH R37GM057260 (A.S.A.). A.S.A is an investigator of the Howard Hughes Medical Institute and the Stowers Institute for Medical Research.

---

**Supplemental Figure 1.**
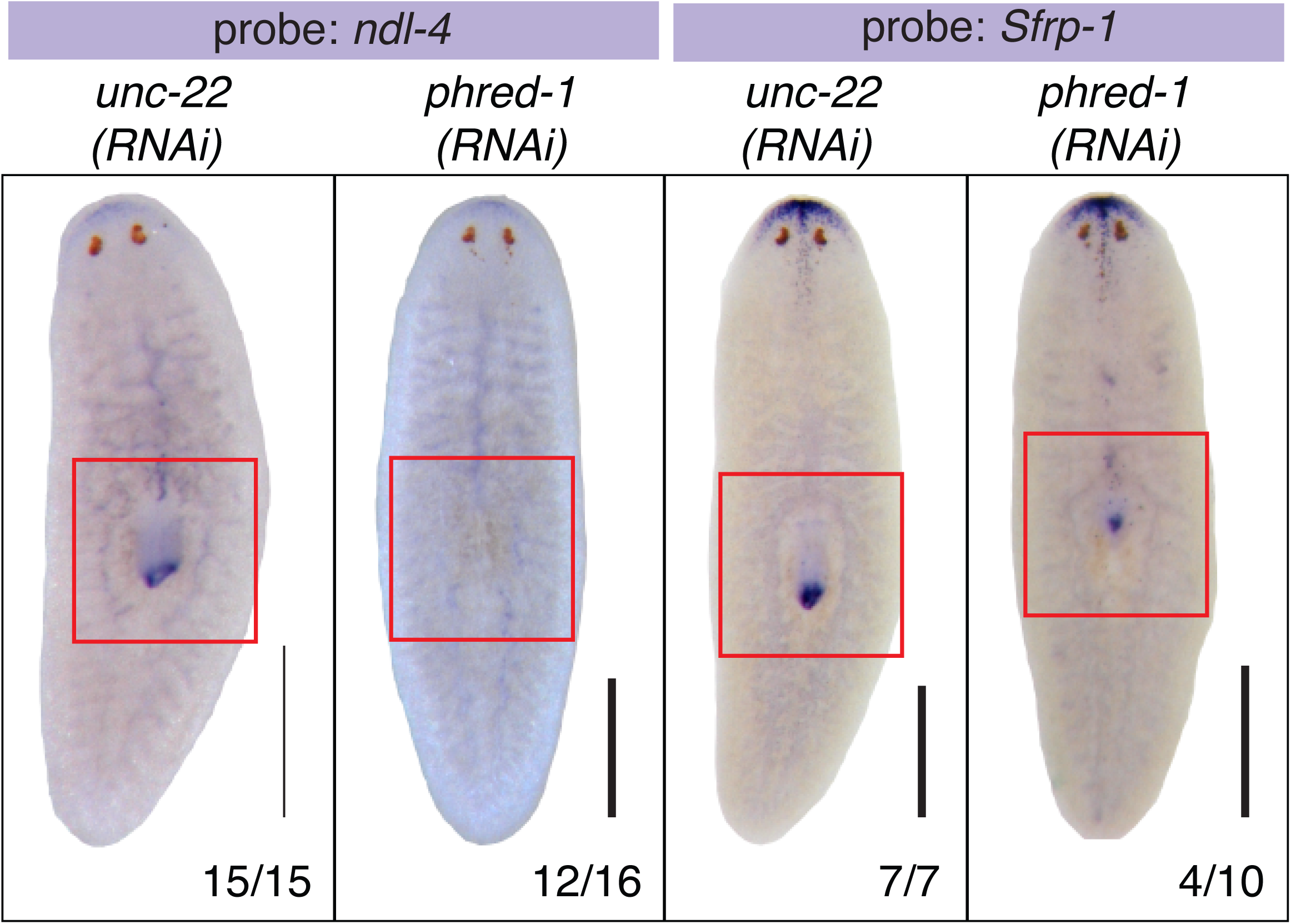
PHRED-1 is required for specification of the distal pharynx. *In situ* hybridization of RNAi animals 7 days after amputation. (A) *ndl-4*, and (B) *Sfrp-1*. N values shown at bottom, from two independent experiments. Scale bars, 500μm.

**Supplemental Figure 2.**
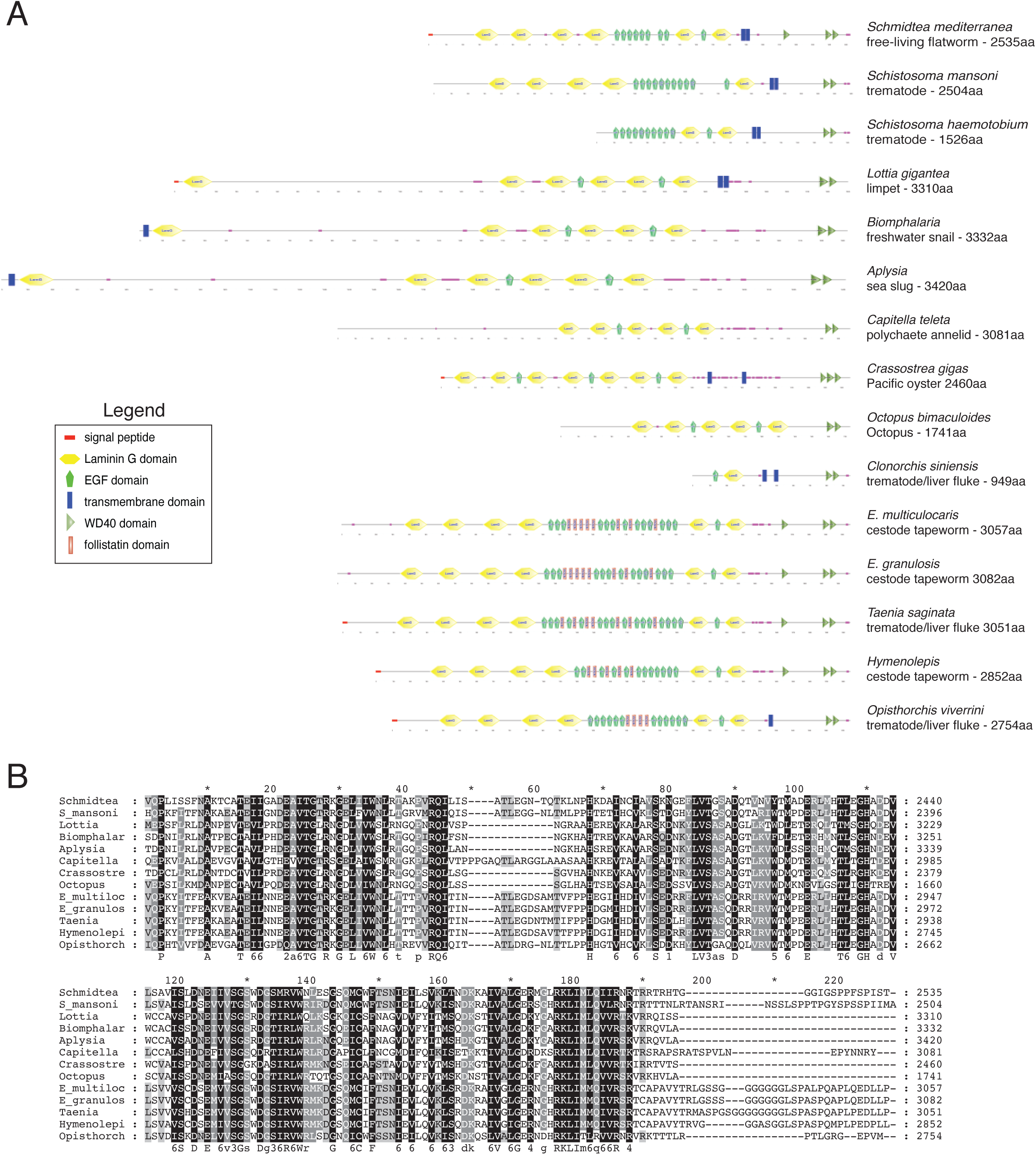
Homology analysis of the PHRED-1 family. (A) BLAST identified 14 homologs of PHRED-1 containing both Laminin G, EGF, and WD-40 domains. Domain predictions are derived from SMART. Homologs from *Clonorchis siniensis* and *Schistosoma haemotobium* are likely incomplete transcripts. The parasitic flatworms *Echinococcus multilocularis, Echinococcus granulosus, Taenia saginata, Hymenolepis microstoma, and Opisthorchis viverrini* have added cohorts of follistatin domains into their extracellular regions. (B) Amino acid alignment of C-terminal WD40 domains of all homologs (excluding incomplete transcripts from *Clonorchis siniensis* and *Schistosoma haemotobium*).

**Supplemental Figure 3.**
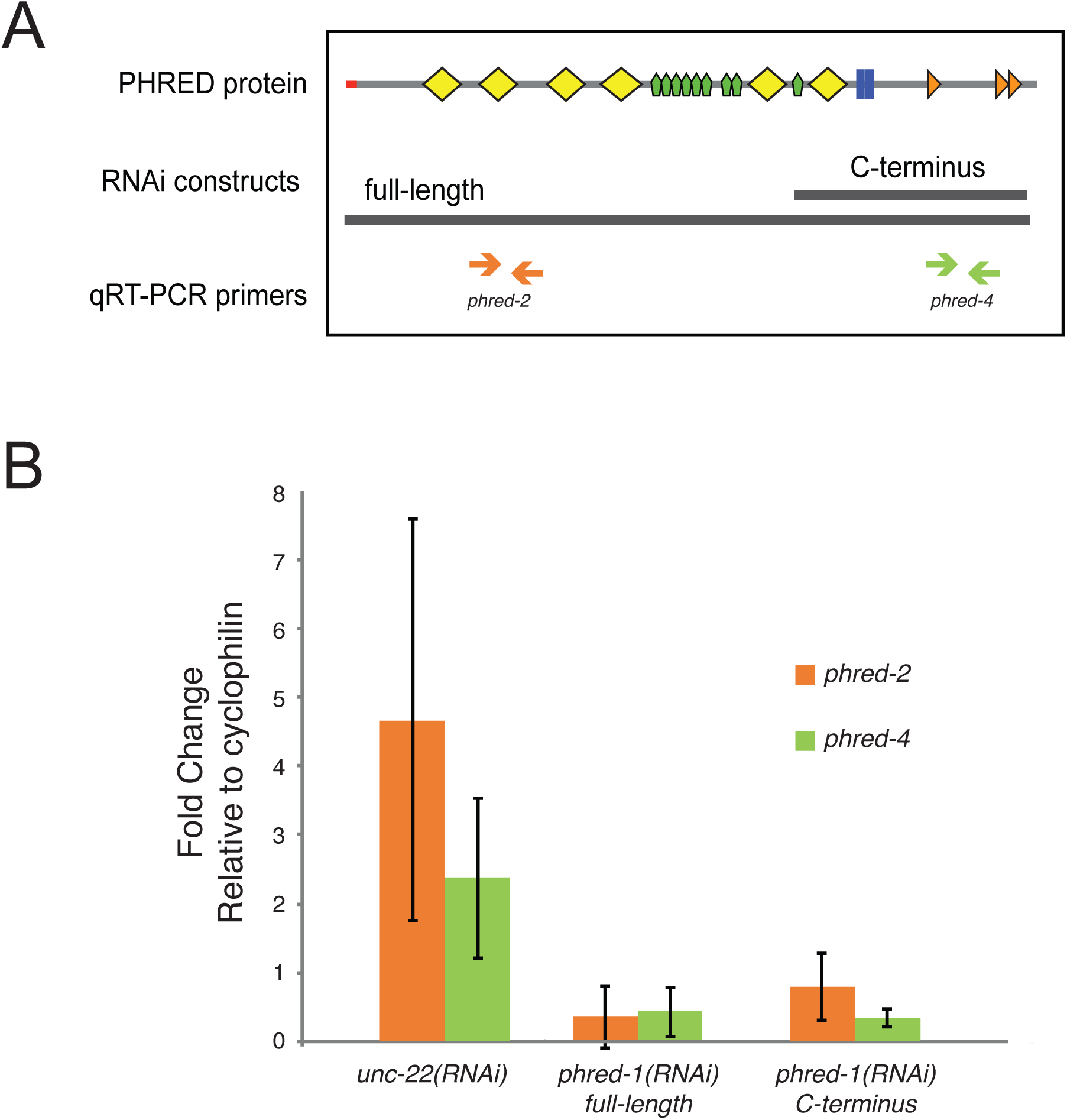
qRT-PCR analysis of *phred-1(RNAi)* knockdown. Top, schematic of PHRED-1 protein showing relative positions of RNAi constructs and qRT-PCR primers used. Bottom, qRT-PCR with indicated primers on *unc-22(RNAi)* or *phred-1(RNAi)* animals induced with different constructs shown above.

**Supplemental Figure 4.**
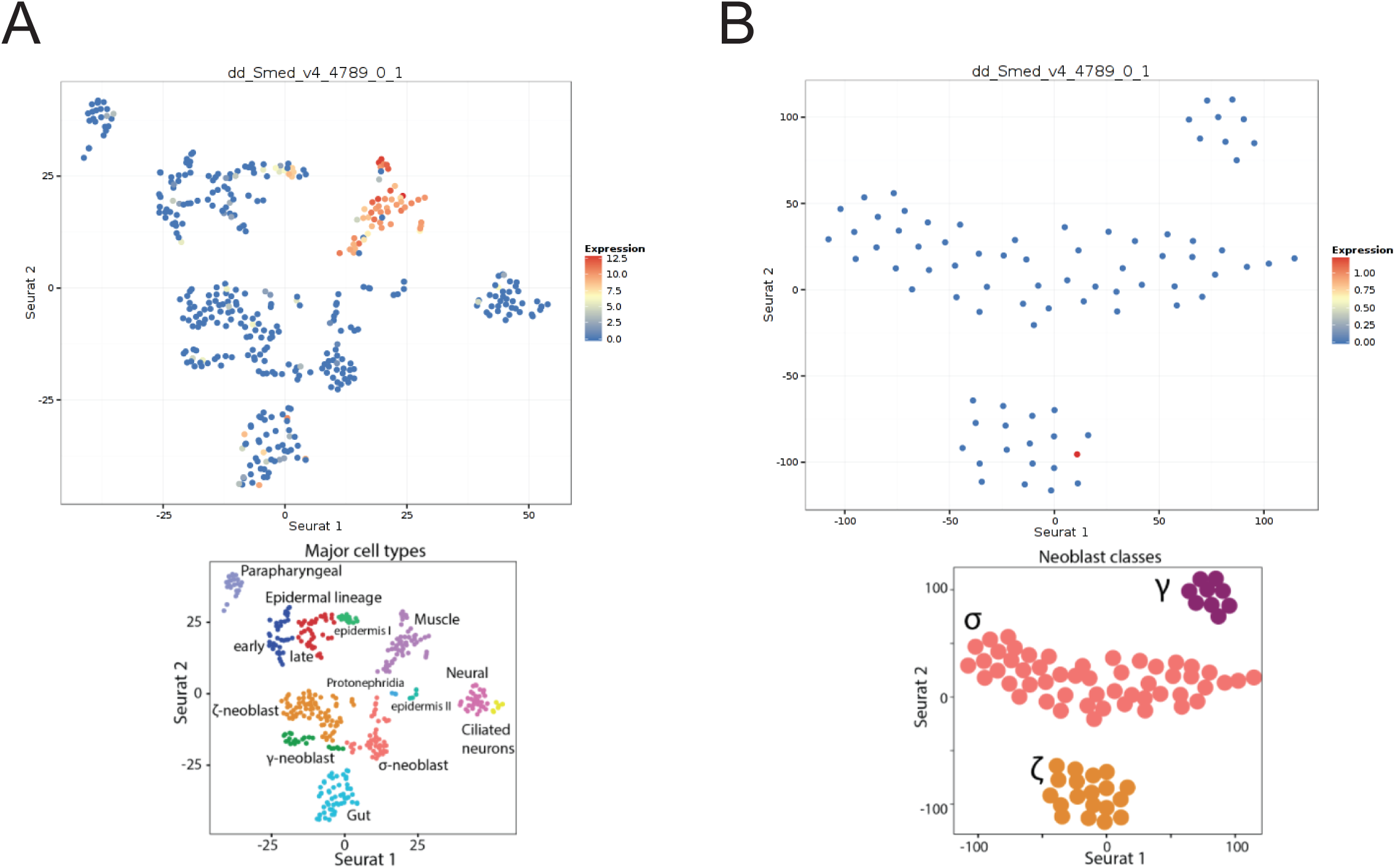
*phred-1* is expressed in muscle cells. (A) Single-cell RNA-sequencing data from (Wurtzel et al., 2015) places *phred-1* (dd_Smed_v4_4789_0_1) among muscle-expressed genes. (B) Sub-categories of neoblasts. *phred-1* is weakly represented among gamma-class neoblasts.

**Supplemental Figure 5.**
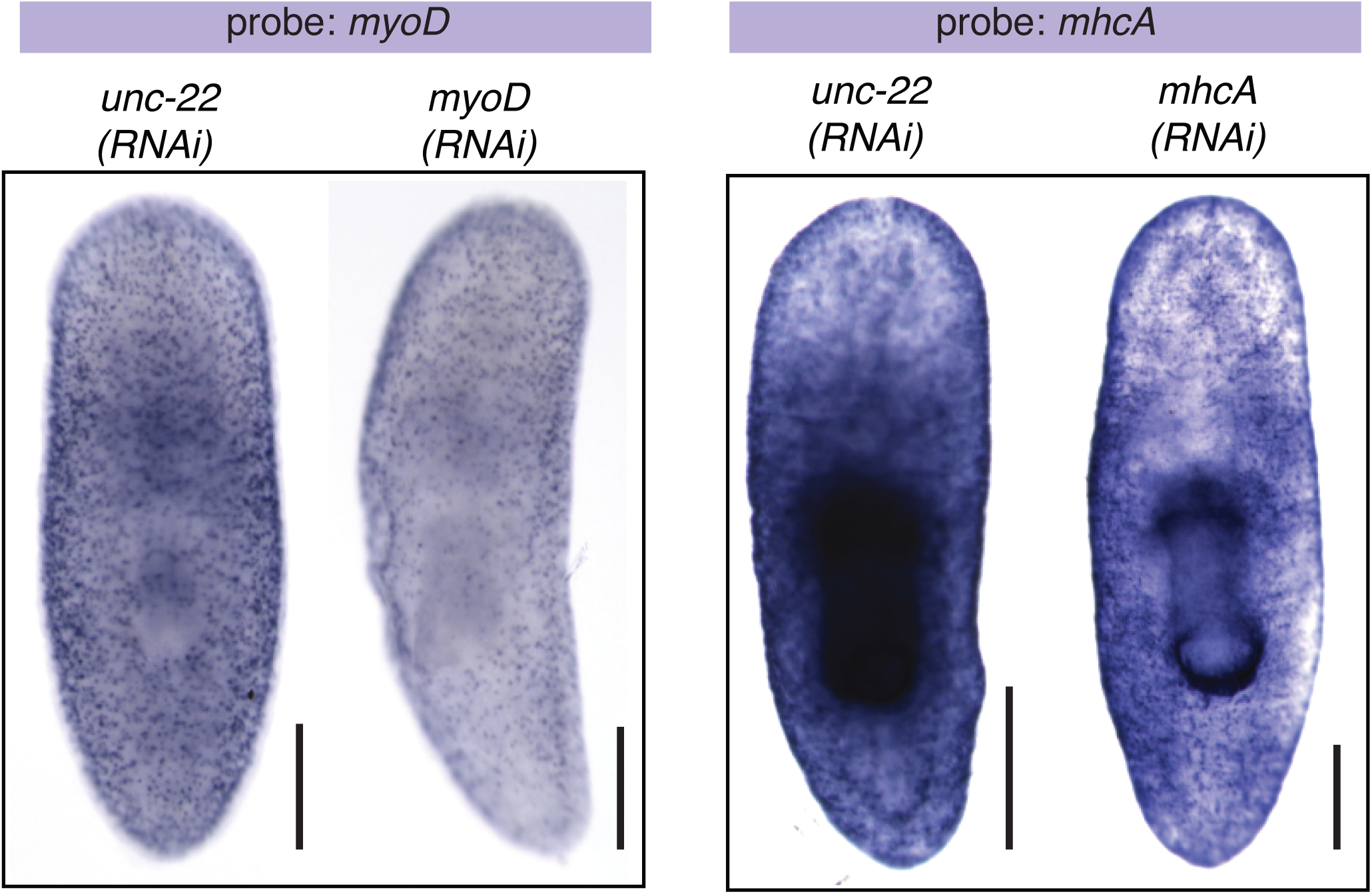
MyoD and mhcA expression after knockdown. *myoD* and *MHCA in situ* hybridizations after RNAi knockdown as indicated. Scale bars, 250μm.

**Supplemental Figure 6.**
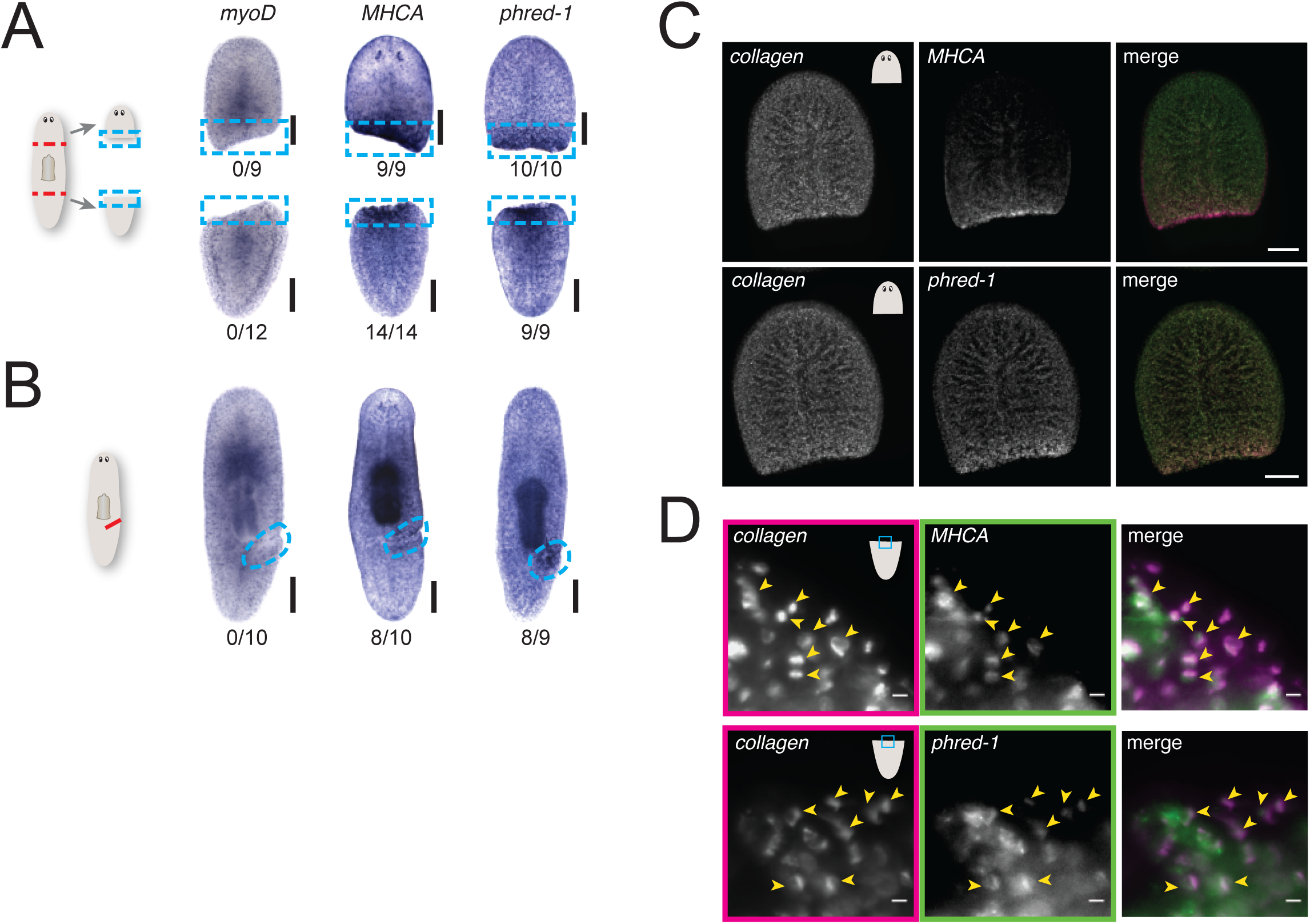
*MyoD* and *collagen* expression are not induced by wounding. (A) Schematic of amputation (dashed red lines). *myoD, MHCA, phred-1 in situ* hybridization 12 hours after amputation. Blue boxes highlight blastema region. (B) Schematic of incision (red line). *myoD, MHCA, phred-1 in situ* hybridization 12 hours after amputation. Blue circles highlight region of incision. (C) Double fluorescent *in situ* hybridization for *collagen* together with *mhcA* or *phred-1* in head fragments 12 hours after amputation. (D) Colocalization of *collagen* with *mhcA* or *phred-1* as in (C), in region highlighted by blue box. Yellow arrowheads highlight colocalization. Scale bars, (A-C) 250μm; (D) 10μm.

